# SFARI Genes and where to find them; classification modelling to identify genes associated with Autism Spectrum Disorder from RNA-seq data

**DOI:** 10.1101/2021.01.29.428754

**Authors:** Magdalena Navarro, T Ian Simpson

## Abstract

**Motivation:** Autism spectrum disorder (ASD) has a strong, yet heterogeneous, genetic component. Among the various methods that are being developed to help reveal the underlying molecular aetiology of the disease, one that is gaining popularity is the combination of gene expression and clinical genetic data. For ASD, the SFARI-gene database comprises lists of curated genes in which presumed causative mutations have been identified in patients. In order to predict novel candidate SFARI-genes we built classification models combining differential gene expression data for ASD patients and unaffected individuals with a gene’s status in the SFARI-gene list.

**Results:** SFARI-genes were not found to be significantly associated with differential gene expression patterns, nor were they enriched in gene co-expression network modules that had a strong correlation with ASD diagnosis. However, network analysis and machine learning models that incorporate information from the whole gene co-expression network were able to predict novel candidate genes that share features of existing SFARI genes and have support for roles in ASD in the literature. We found a statistically significant bias related to the absolute level of gene expression for existing SFARI genes and their scores. It is essential that this bias be taken into account when studies interpret ASD gene expression data at gene, module and whole-network levels.

**Availability:** Source code is available from GitHub (https://doi.org/10.5281/zenodo.4463693) and the accompanying data from The University of Edinburgh DataStore (https://doi.org/10.7488/ds/2980)

## 1 Introduction

Autism spectrum disorder (ASD) encompasses a diverse group of developmental disorders characterised by deficits in social interaction, impaired communication skills, and a range of stereotyped and repetitive behaviours (Lord *et al*., 1994). ASD has a strong genetic component, with heritability estimated to be as high as 52% (Gaugler *et al*., 2014) and hundreds of genes believed to be disrupted by it (Iossifov *et al*., 2014), however, for 75% of the cases, the causes still remain unknown (Quesnel-Vallières *et al*., 2018), which suggests there is still a lot to discover about this complex and heterogeneous disorder.

There are many approaches to study the genetic components underlying the aetiology of ASD. The most direct, and one of the most popular approaches, is to study likely causative mutations that have been found in patients with the disorder. Arguably the largest source of these are the Simons Foundation Autism Research Initiative (SFARI) (Banerjee-Basu and Packer, 2010) who created SFARI-gene, a constantly evolving, expertly curated database of candidate genes involved in autism susceptibility by integrating genetic information from multiple research studies. It currently consists of 992 genes, which have been scored with a value from 1 to 3 reflecting the strength of the evidence linking a gene to ASD, where a score of 1 is assigned to genes that have a high confidence of being implicated, 2 to strong candidates, and 3 to genes that only have relatively weak evidence supporting their connection to ASD.

Another common approach is to compare gene expression between ASD patients and unaffected controls using transcriptomics. This has led to the discovery of many candidate genes for ASD and identified convergent molecular processes involved in the disorder (Pinto *et al*., 2014). These analyses have also revealed interactions between molecular pathways and other contributory factors and have helped us to understand how diverse mechanisms and risk factors can combine to produce complex behavioural outcomes (Quesnel-Vallières *et al*., 2018).

Despite the differences between the SFARI-gene curated list and transcriptomic data, they are frequently used together, either using genes from SFARI-gene to design transcriptomic experiments or to validate results (Araujo *et al*., 2017; Berto *et al*., 2018; Gokoolparsadh *et al*., 2017; Lombardo *et al*., 2017; Nowakowski *et al*., 2017; Yu and He, 2017; Suetterlin *et al*., 2018; Wang *et al*., 2018). More recently, studies have begun to combine information from these two sources into single models that learn jointly from these data (Brueggeman *et al*., 2020; Cogill and Wang, 2016; Di Nanni *et al*., 2019; Lin *et al*., 2020).

We aim to study how best to use these data together; focusing on when it is appropriate to combine them and what aspects should be taken into consideration when doing so. We do this in three ways: at the *gene-level*, by examining individual genes independently from one another, at the *module-level*, by examining genes in groups defined by similarities in gene expression profiles, and finally at the *systems-level*, analysing all of the genes simultaneous in a fully-connected expression network.

## 2 Methods

Pre-processing and analysis of transcriptomic data was performed using the DESeq2 (Love *et al*., 2014) and WGCNA (Langfelder and Horvath, 2012) software packages. We developed the classification models using features from the gene co-expression analysis. Details for these steps are described in the following sections and the full data and code are freely available through the links provided in the Abstract.

### 2.1 Datasets

The version of the SFARI Gene dataset used corresponds to Q1 2020. It contains 1114 genes, of which 202 genes have a score of 1, 239 a score of 2 and 586 a score of 3. The 87 genes that weren’t assigned a score were not included in the analysis.

For the transcriptomic data, three RNA-seq datasets were studied, all consisting of human post-mortem brain tissue samples belonging to ASD individuals as well as a non-psychiatric control group. The main dataset was obtained from the GitHub repository from (Gandal *et al*., 2018). It contains 88 samples; 53 belonging to 24 ASD individuals and 35 to 17 controls, corresponding to the frontal, temporal, parietal and occipital cortical regions. After preprocessing, the final dataset contains 16132 genes and 80 samples. The first supporting dataset corresponds to (Gupta *et al*., 2014). It contains 104 samples; 47 belonging to 32 ASD individuals and 57 to 40 controls, extracted from Brodmann Areas 10, 19 and 44. The final version of this dataset contains 13162 genes and 89 samples. And the second supporting dataset was obtained from (Wright *et al*., 2017), the expression matrix was downloaded from (Zoubarev *et al*., 2012) and the metadata information from NCBI’s Gene Expression Omnibus (Edgar and Lash, 2002) with Series accession number GSE102741. It contains 52 samples, all corresponding to the dorsolateral prefrontal cortex; 13 of these belong to ASD individuals and 39 to controls. The final version of this dataset contains 15392 genes and 49 samples.

To broadly define genes that had neuronal functions we annotated genes using Gene Ontology annotations (Ashburner *et al*., 2000; Consortium and Acencio, 2018) if their term name or description contained the substring ‘neuron’. All comparisons performed between SFARI genes and other genes within the gene expression data are performed separately, allowing us to compare SFARI genes to neuronal, non-neuronal and non-SFARI genes as required.

“Krishnan-scores” were obtained from genome-wide autism-gene predictions available from http://asd.princeton.edu as part of the supplementary material from (Krishnan *et al*., 2016). “TADA-scores” were extracted from Table S3 in (He *et al*., 2013), and “DisGeNET-scores” were retrieved using the “disgenet2r” R package (Piñero *et al*., 2019).

### 2.2 Data Preprocessing

Meta-data for genes were retrieved from NCBI (Sayers *et al*., 2011) using the bioMart package (Durinck *et al*., 2005). During filtering we retained known protein coding genes. Of these, genes with a high percentage of zeroes across all samples were removed. The threshold for this was determined as the minimum percentage of zeroes where the strongest heteroscedasticity patterns in the normalised dataset disappears. We next removed outlier samples by calculating the pairwise correlation between expression profiles, then aggregating these for each sample and calculating their distance to the rest of the samples as a group. Outlier samples were identified if this distance was larger than two standard deviations away from the mean.

For Differential Expression Analysis (DEA), first, the SVA package (Leek *et al*., 2019) was used to calculate the surrogate variables associated with unknown sources of batch effects in the data, and then, the DESeq2 package was used to perform DEA, using Diagnosis as target and including the batch-related features as well as the surrogate variables obtained from SVA into the formula. The null hypothesis used for the analysis was a log fold change threshold of 0. After this, the data was normalised using the *vst* function from the DESeq2 package. Finally, batch effects were corrected for using a linear transformation to remove the effects captured by the surrogate variables from the *SVA* and *ComBat* functions removing the batch effects captured by the original features of the samples.

After preprocessing, the main feature that characterises our samples is their Diagnosis status. This achieved perfect separation of the samples using only the first principal component (Figure 1).

**Fig. 1.**
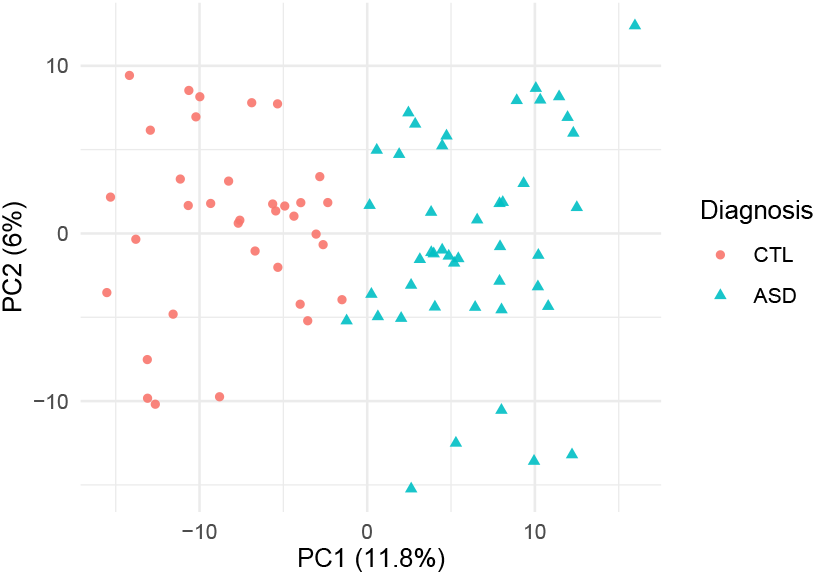
PCA plot of the samples in our main transcriptomic dataset characterised by their expression profiles after preprocessing. The numbers in parenthesis on the axis represent the percentage of variance explained by each component.

### 2.3 WGCNA and Enrichment Analysis

The network was built using the Weighted Gene Correlation Network Analysis (WGCNA) package. The expression matrix was transformed using the biweight midcorrelation metric, with the *signed hybrid* and *pickSoftThreshold* functions to obtain a scale-free topology, and subtracting the resulting topological overlap matrix from 1.0 to obtain the dissimilarity matrix. Clusters within this matrix were identified using hierarchical clustering with the *cutreeDynamic* algorithm. The strength of the relation between each of these modules and Diagnosis status was measured with the correlation of the module’s first principal component (Eigengene) and the Diagnosis feature vector of the samples belonging to that module. Modules with a correlation magnitude higher than 0.9 were considered to have a strong correlation with Diagnosis status.

The enrichment in SFARI genes within a module was calculated using the Over Representation Analysis (ORA) provided by the clusterProfiler package (Yu *et al*., 2012); the modules with a bonferroni corrected p-value lower than 0.05 were labelled as having a statistically significant enrichment in SFARI genes. The Enrichment Analysis for each module was performed using the Gene Set Enrichment Analysis (GSEA) as well as the ORA in the KEGG (Kanehisa and Goto, 2000), Reactome (Croft *et al*., 2013), Gene Ontology, Disease Ontology (Schriml *et al*., 2018) and Disease Gene Network datasets (Piñero *et al*., 2019), using a bonferroni corrected p-value of 0.1 on each of these two enrichment analyses to identify candidate enriched terms, and labelling the terms that were found to be candidates in both methods as enriched in the terms.

### 2.4 Classification Model

The dataset used to train the classification model consists of all the genes that were assigned to a module by WGCNA, characterised by a set of descriptive variables and a binary objective variable indicating if the gene is included in the SFARI-gene set or not, ignoring the SFARI scores. The descriptive variables selected for the model are the correlation of a gene’s expression pattern to Diagnosis status (Gene Significance), including both the original correlation and its absolute value; the correlation of a gene’s assigned module to Diagnosis status (Module-Trait correlation); and the gene’s correlation to the Eigengene of each of the modules in the network (Module Membership). The resulting dataset consists of 15,994 observations, 58 descriptive variables, and one objective variable, which contains 789 positive and 15,211 negative values.

The genes are separated into training and testing sets, using 75% of the genes in the training set, where the imbalance between labels is corrected using the SMOTE over-sampling technique (Chawla *et al*., 2002), the remaining 25% of the genes are reserved as a test set.

Ridge regression (Hoerl and Kennard, 2000) was selected as the classification model because of the strong multicollinearity found in the descriptive variables in the dataset, using repeated cross validation to estimate the optimal value for the regularisation parameter of the model using 10-fold cross validation with 5 repeats. The model is trained 100 times using different partitions of the training and testing sets and the results from each of the runs are combined for the calculation of the final predictions and performance evaluation of the model. The performance metrics used are Area Under the ROC Curve (AUC), which measures the ability of a classifier to distinguish between classes; the Maximum Lift Point (MLP), which reflects the density of positive observations (SFARI genes) in the observations with the highest assigned probabilities in the dataset; and the Balanced Accuracy, which is a commonly used substitute for the regular Accuracy metric when the classes are imbalanced, and is the average of the proportion of positive observations that were correctly classified, and the proportion of negative observations that were correctly classified.

As a modification to this original regression model, the weighting technique proposed by (Jiang and Nachum, 2019) was used to correct the bias found in the model related to the mean level of expression of the genes, using Demographic Parity as a measure for bias, and modifying the constraint function to measure not only if there is bias or not, but the magnitude of this bias:

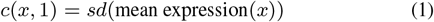

Where *x* is each of the genes that are labelled as 1 by the model.

The top candidate gene list comprises those genes with the highest probabilities assigned by the final model and represent genes that share features in common with existing SFARI-genes. To allow calculation of the standard deviation of the performance metrics, the whole model, including the repetitions for different training-testing partitions, was repeated 100 times.

## 3 Results

### 3.1 Mean Level of Expression

Before starting the analysis of the SFARI genes, it is important to notice the significance of the role the mean expression of the genes play in a transcriptomic dataset. This value is calculated averaging the level of expression of each gene across all samples, regardless of diagnosis. As can be seen in Figure 2, genes are strongly characterised by their mean level of expression, explaining over 99% of the total variance in the dataset.

**Fig. 2.**
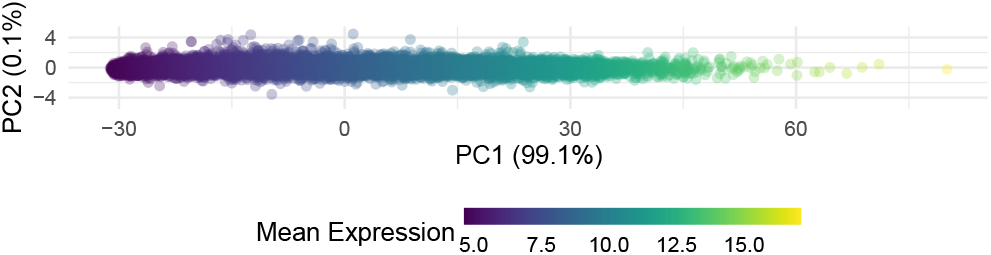
PCA plot of the genes characterised by their expression patterns across all samples and coloured by their mean level of expression. The x-axis corresponds to the first principal component, which explains over 99% of the variance, is strongly related to the level of expression of the genes

Comparing the mean level of expression of the genes that correspond to SFARI against the rest of the genes in our transcriptomic dataset, we can see that they have a statistically significant higher level of expression to both of the other gene groups with a Benjamini-Hochberg corrected p-value lower than 10^-4^, as it can be seen in Figure 3(a), agreeing with the results presented in (Lin *et al*., 2020).

**Fig. 3.**
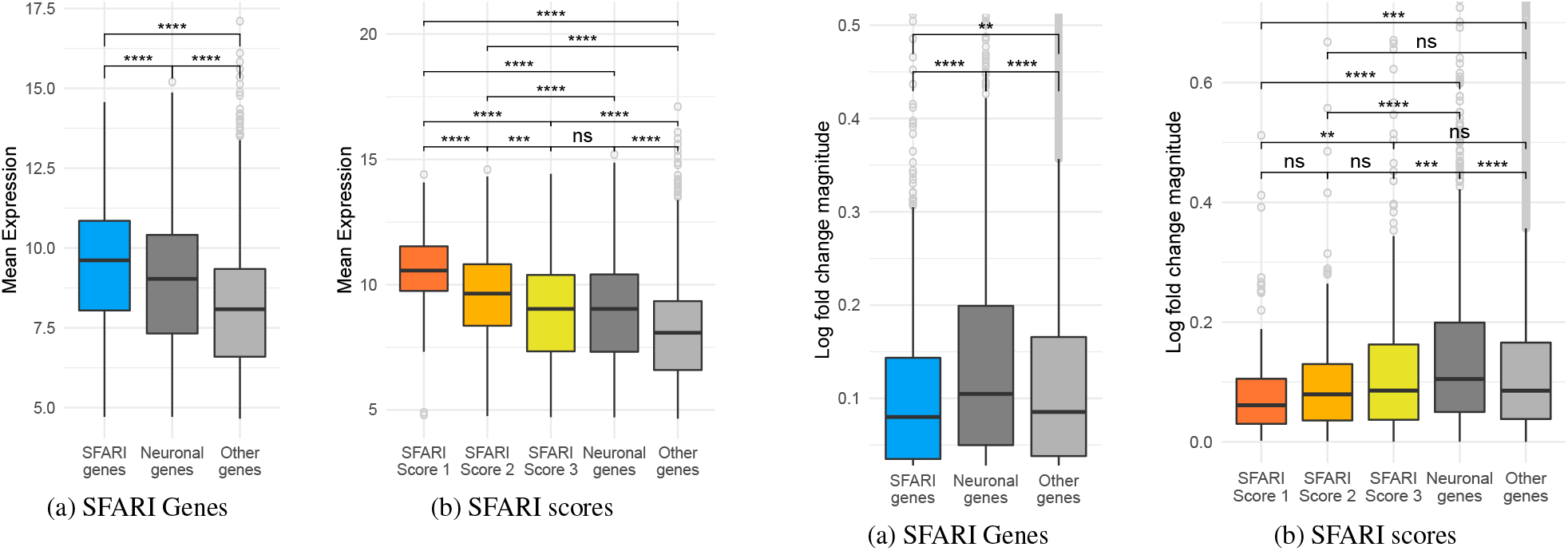
Box plot of the mean level of expression of the genes, comparing the SFARI genes against genes with neuronal annotations as we as with the rest of the genes in the dataset. The brackets at the top indicate pairwise comparisons, using a Welch t-test to study if the differences in level of expression between groups are statistically significant, and the asterisks indicate the magnitude of the corrected p-value of each test: ns = p-value>=0.05, * = p-value < 0.5, ** = p-value < 0.01, *** = p-value < 0.001, and **** = p-value < 0.0001

Figure 3(b) shows that separating the SFARI Genes by SFARI scores, we find a similar pattern; the higher the SFARI score, the higher the level of expression of the genes, with genes belonging to SFARI score 1 having the highest level of expression of all groups, followed by SFARI score 2 and then by SFARI score 3. All of the differences between groups are statistically significant with a corrected p-value lower than 10^-3^, even between SFARI scores, except for the comparison between SFARI score 3 and the neuronal genes, where the difference is not statistically significant.

### 3.2 Gene Level

In order to determine the relationship between SFARI Genes and their differential expression between ASD and control groups we investigated how the log fold change magnitude varied with the percentage of differentially expressed genes by category. We find that SFARI genes have a consistently lower percentage of differentially expressed genes when compared to the neuronal group, and very similar values to the rest of the genes, regardless of the log fold change threshold (Figure 4). All categories of SFARI score have very similar percentages of differentially expressed genes, so only the combined is shown here for clarity.

**Fig. 4.**
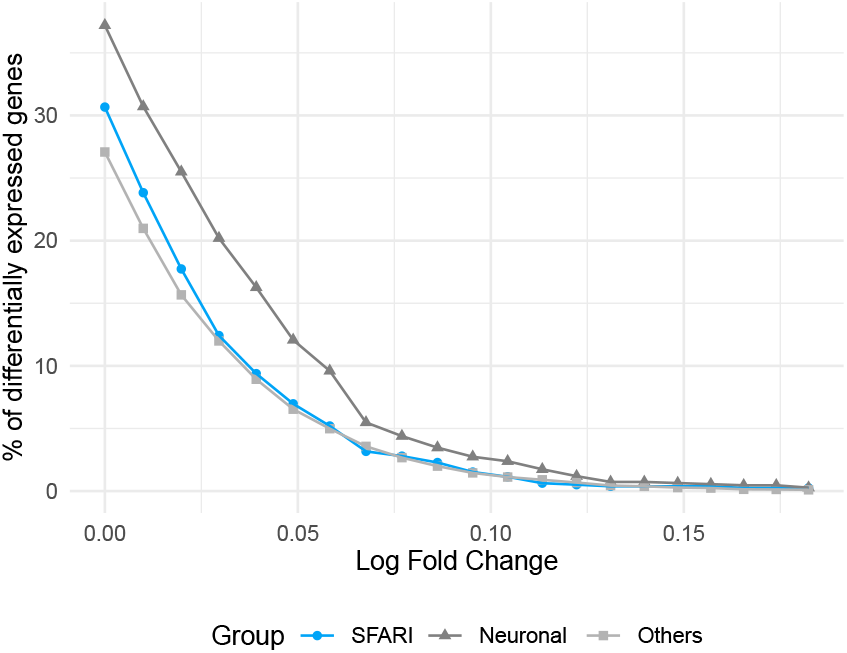
Percentage of genes found to be differentially expressed using different log fold change thresholds

If we now compare the log fold-change magnitude of genes in each category we find that the SFARI genes have statistically significantly lower values than both neuronal genes and other genes, with a corrected p-value lower than 0.01 (Figure 5(a)).

**Fig. 5.**
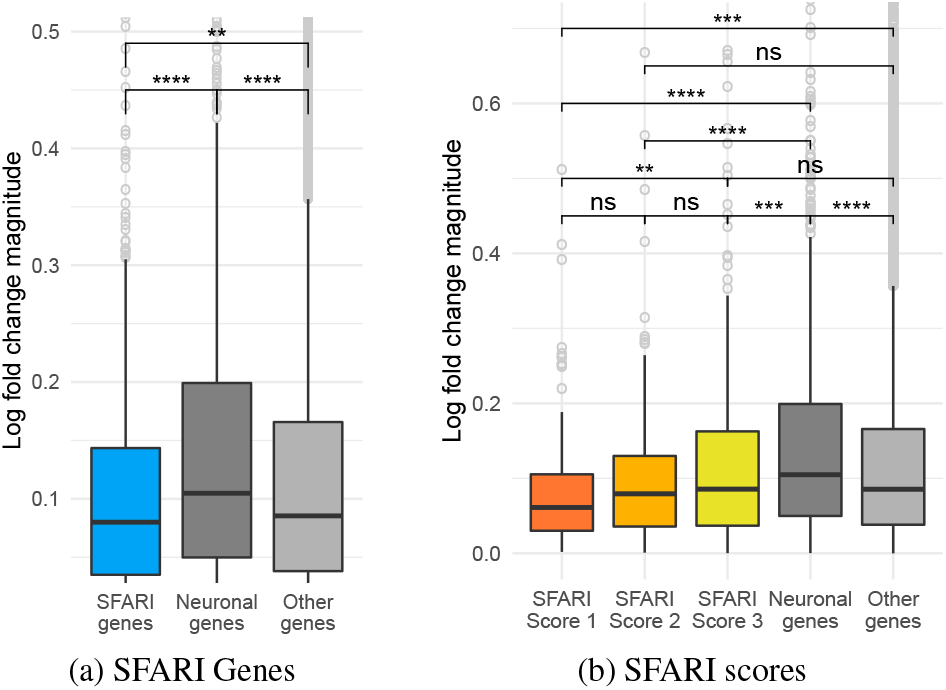
Box plot of the log fold change magnitude of the genes, comparing the SFARI genes against genes with neuronal annotations and with the rest of the genes in the dataset. As before, the asterisks at the top indicate the magnitude of the corrected p-value from pairwise Welch t-test comparisons to study if the differences between groups is statistically significant.

Separating the SFARI-genes by SFARI scores we find that the higher the SFARI score, the lower the log fold change magnitude, with SFARI score 1 having the lowest values of all groups, including the rest of the genes that are neither SFARI nor neuronal, and neuronal genes having the highest. These differences are statistically significant for all other groups (p < 10^-3^). The differences between adjacent SFARI scores, as well as between the group of non-SFARI, non-neuronal genes and SFARI scores 2 and 3 were the only was that were not found to be statistically significant.

### 3.3 Module Level

We next explored the relationship between the correlation of WGCNA’s modules with diagnosis status and the distribution of SFARI-genes to determine whether SFARI-genes are strongly associated with those modules that have strong correlation with ASD.

The WGCNA package finds 55 modules within the co-expression network, leaving only 138 genes unassigned to any module. To compare the influence the diagnosis of the samples has in each module against the presence of SFARI genes, we use the Module-Trait correlation, and 1 minus the p-value of the Over Representation Analysis of the SFARI genes, respectively (Figure 6), where each point represents a module, and the axes corresponds to each of these two characteristics. The relatively flat trend line in the plot, as well as what seems a relatively uniform distribution of the modules statistically significantly enriched in SFARI genes across the different levels of module-diagnosis correlation suggest there may not be a strong relation between them. The most notable change in the trend line is a decrease in the enrichment in SFARI Genes for the group of genes with the highest positive module-diagnosis correlations, and perhaps a small increase in statistically significantly enriched modules in SFARI genes in the modules with the most negative module-diagnosis correlations, but neither of these patterns is particularly strong.

**Fig. 6.**
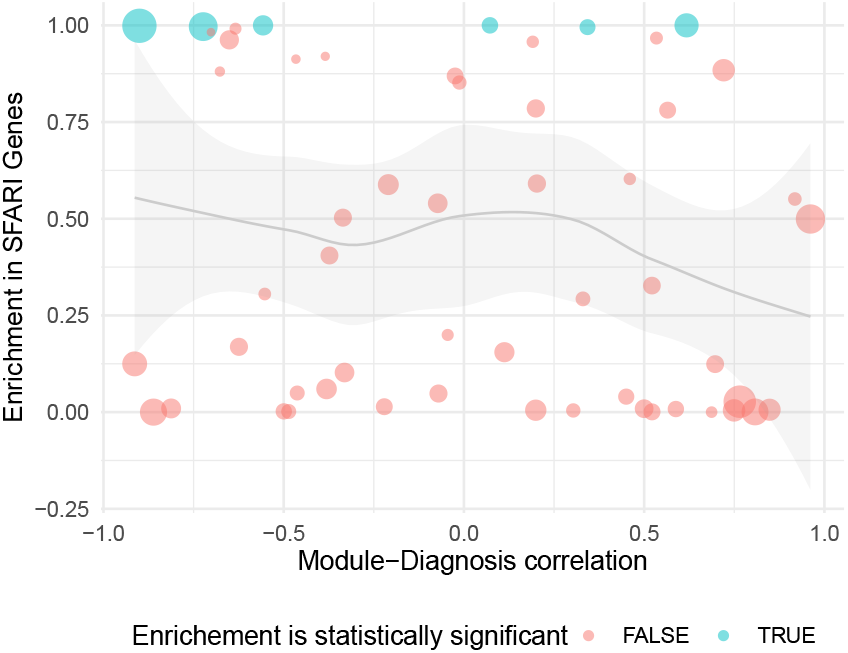
Scatter plot of the WGCNA modules comparing the strength of the correlation of the modules to the diagnosis of the samples and the enrichment of SFARI Genes. Each point represents a module; its position on the plane is defined by these two metrics, its size corresponds to the number of genes in the module, and its colour indicates if the enrichment in SFARI scores was statistically significant. The grey line corresponds to the trend line illustrating the relation between the two variables we are studying, with the shaded area around the line displaying its 95% confidence interval.

Performing a similar analysis by substituting the module-diagnosis correlation of each module for the mean level of expression of the genes it contains, we get a much clearer pattern: as Figure 7 shows, modules with higher levels of expression have a higher enrichment in SFARI genes, and none of the modules where the enrichment in SFARI genes was found to be statistically significant have a low mean level of expression. These results are consistent with the findings presented in section 3.1, and show that the positive relation between level of expression and SFARI genes persist at module level.

**Fig. 7.**
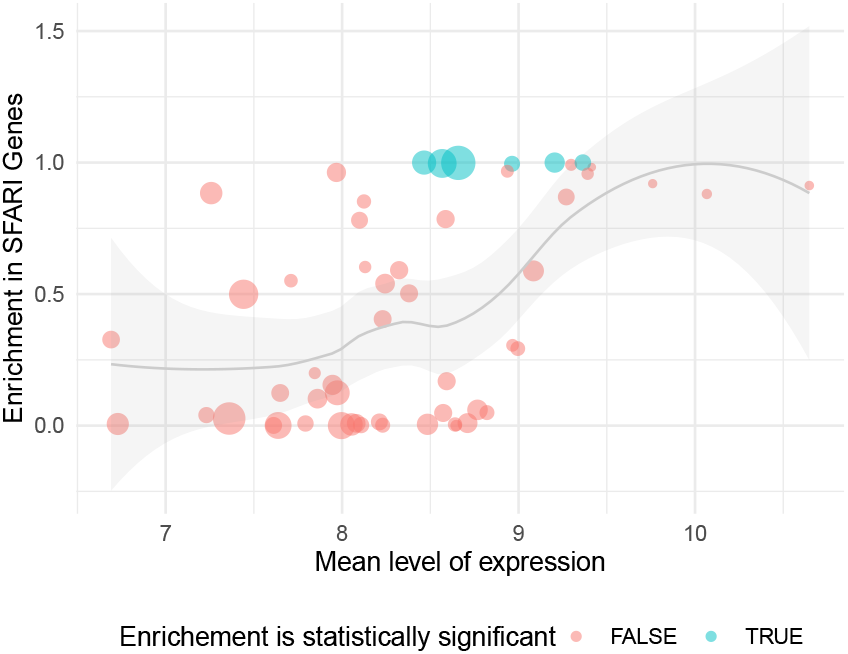
Scatter plot of the WGCNA modules this time comparing the mean level of expression of the genes contained in each module against its enrichment in SFARI Genes. The details of the plot are the same as in Figure 6

### 3.4 Systems Level

We next built a classifier to assign a probability to non-SFARI genes based on their similarity to SFARI-gene differential expression profiles between ASD and control samples and features extracted from our coexpression network modules. By measuring the performance of the model we quantified the reliability of the model and for the genes with the highest probabilities evaluated whether biological evidence existed in the literature to support their relevance to ASD.

Table 1 shows the performance of the two models described in Section 2.4 and compares them to a third model, where the classifier used is the same, but the SFARI labels in the classification dataset have been shuffled at random, to give a baseline against which we can compare the performance of the other two models.

**Table 1.**
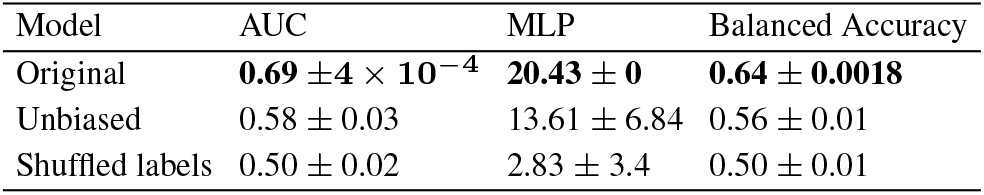
Performance metrics of the two classification models used as well as a third model using a shuffling of the SFARI labels in the data. The highest value for each performance metric is represented in bold.

| Model | AUC | MLP | Balanced Accuracy |
| --- | --- | --- | --- |
| Original | **$0.69 \pm 4 \times 10^{-4}$** | **$20.43 \pm 0$** | **$0.64 \pm 0.0018$** |
| Unbiased | $0.58 \pm 0.03$ | $13.61 \pm 6.84$ | $0.56 \pm 0.01$ |
| Shuffled labels | $0.50 \pm 0.02$ | $2.83 \pm 3.4$ | $0.50 \pm 0.01$ |

The original Ridge regression model performs well, as we can see in Table 1, but when we compare the mean expression of the genes against the probability assigned to them by this model, we find that there is a strong positive relation for all genes except for the ones with the lowest levels of expression, as Figure 8 shows. This suggests that the classifier is using the level of expression of a gene, or some confounder of it, as a factor when calculating its similarity to the SFARI genes, which was expected, since this relation had already been noticed both at gene- and module-level.

**Fig. 8.**
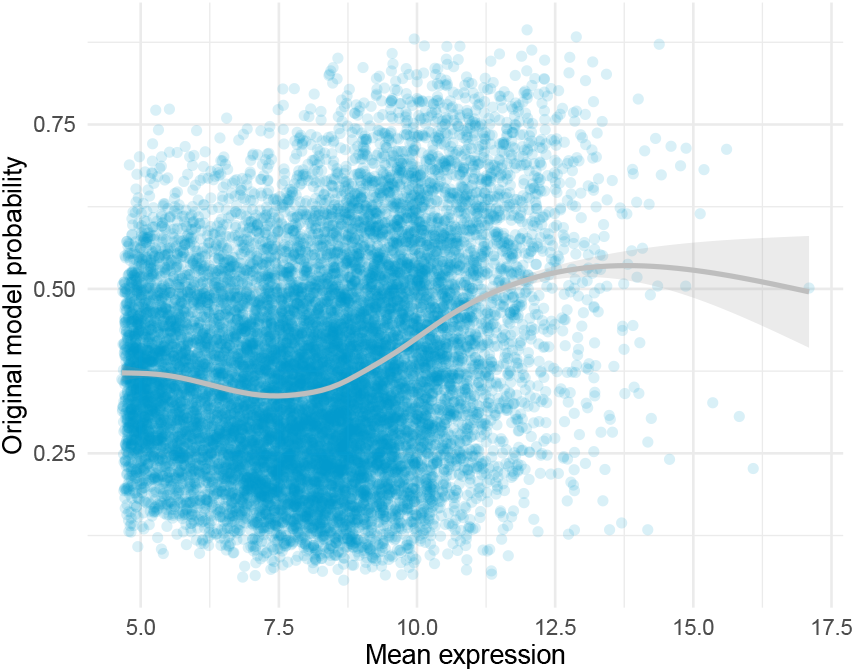
Scatter plot of the genes in the classification dataset, the x-axis corresponds to their mean level of expression and the y-axis to the probability assigned by the model indicating how likely they are to be SFARI genes. The grey line corresponds to the trend line illustrating the relation between these two features, with the shaded area around the line displaying its 95% confidence interval.

We modified the algorithm to correct for the relationship between SFARI genes and mean level of expression using the weighting technique, as detailed in Section 2.4, after which the strongest patterns connecting the mean level of expression and the probability of the model are removed (Figure 9), although it looks like the correction may have been too strict, since the trend line now has a negative slope. This new version of the algorithm, which we call the “unbiased” model, has a worse performance than the original model, as seen in Table 1, because it is no longer using the mean expression of the genes to identify the SFARI genes, but is still performing better than the “shuffled” model.

**Fig. 9.**
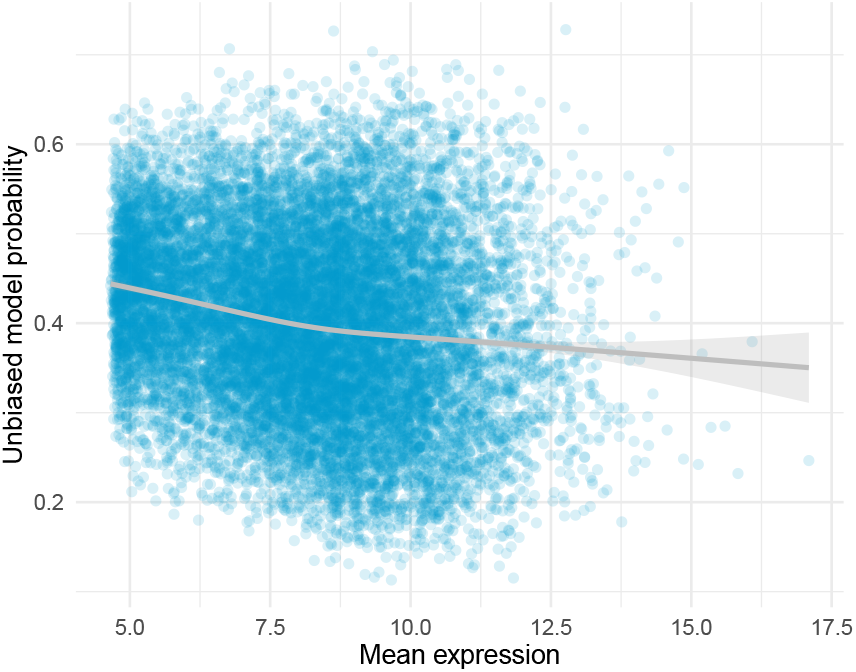
Scatter plot of the relation between the mean level of expression of the genes and the scores from the unbiased classification model.

Table 2 shows the non-SFARI genes that were assigned the highest probabilities by the unbiased model. All of these genes have been found to have some connection with ASD, and gene CORO1A has subsequently been included in the SFARI-gene list with a score of 1. This suggests the model is indeed able to identify genes with similar behaviour to SFARI genes and that the results also have biological relevance to ASD. We find that when a *systems-level* network approach to differential gene expression modelling is combined with categorical labelling of disease genes in a suitably optimised predictive classification model we can successfully identify novel candidate genes that are not found by using either *gene-level* or *module-level* based approaches.

**Table 2.**
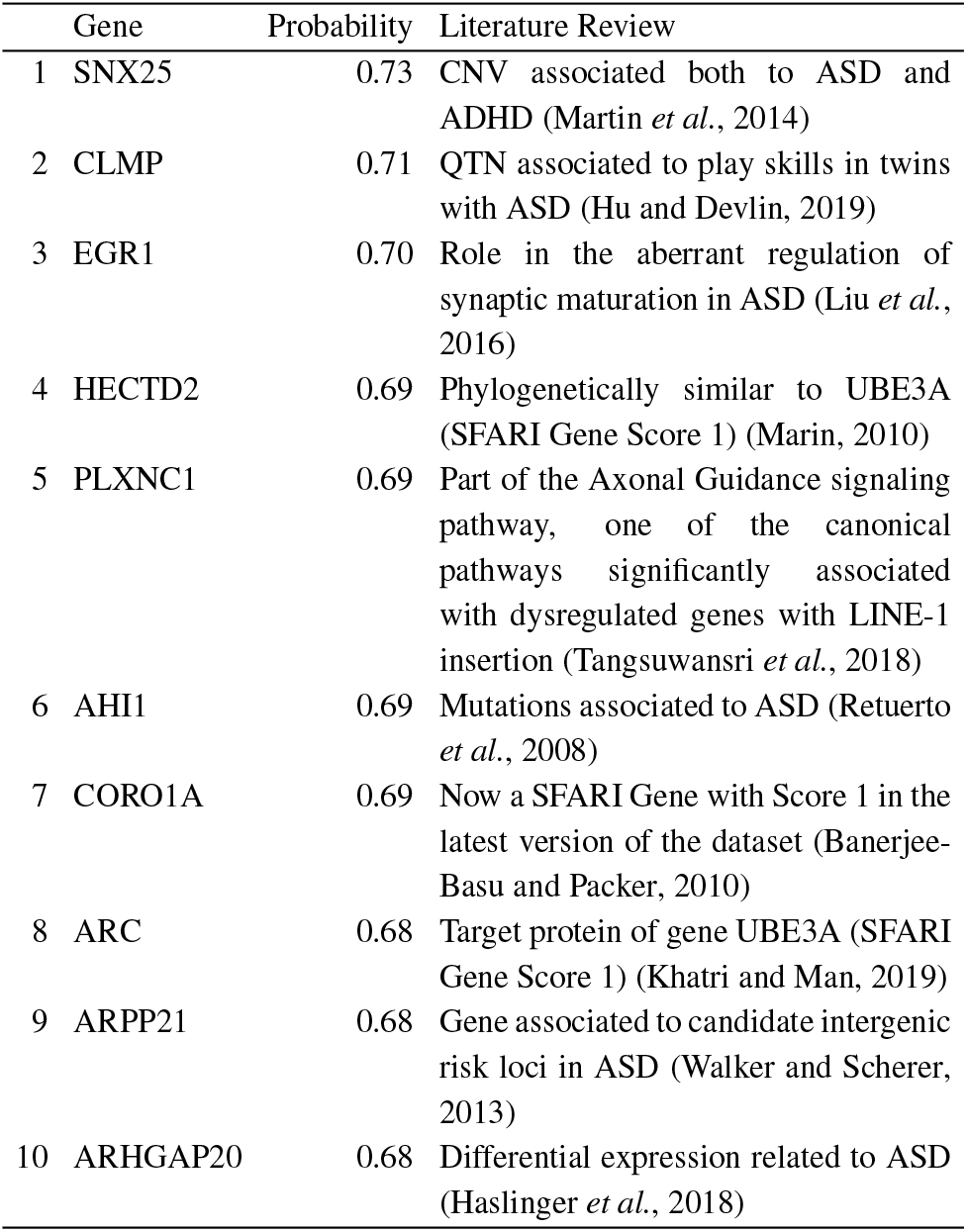
Top 10 non-SFARI genes with the highest probabilities assigned by the unbiased model.

### 3.5 Comparison with other scoring systems and disorders

Given the strong pattern related to the mean level of expression of the genes found in the SFARI Genes dataset, we wanted to know if this pattern was present in other lists of candidate ASD genes, as well as in genes believed to be involved in other neurodevelopmental disorders.

#### 3.5.1 Other ASD scoring systems

Three ASD scoring systems were selected to compare against SFARI: the Krishnan probability score, which uses a gene co-expression network and a list of ASD genes to train a classifier; the Sanders TADA score, which uses whole-exome sequencing to incorporate information from *de novo* mutations, inherited variants present, and variants identified within cases and controls to create a gene-based likelihood model; and the DisGeNET score, which integrates information from various repositories. All of these scores are continuous instead of categorical like SFARI, so we use the Pearson correlation to make pairwise comparisons between these scores and the Polyserial correlation to compare them to the SFARI Genes. As Figure 10 shows, the SFARI, DisGeNET and Krishnan scores have a strong correlation, while Sanders TADA score has either a neutral or a negative correlation with all the others. All of the correlations have a p-value lower than 0.05, the highest being Krishnan vs. TADA with 0.04.

**Fig. 10.**
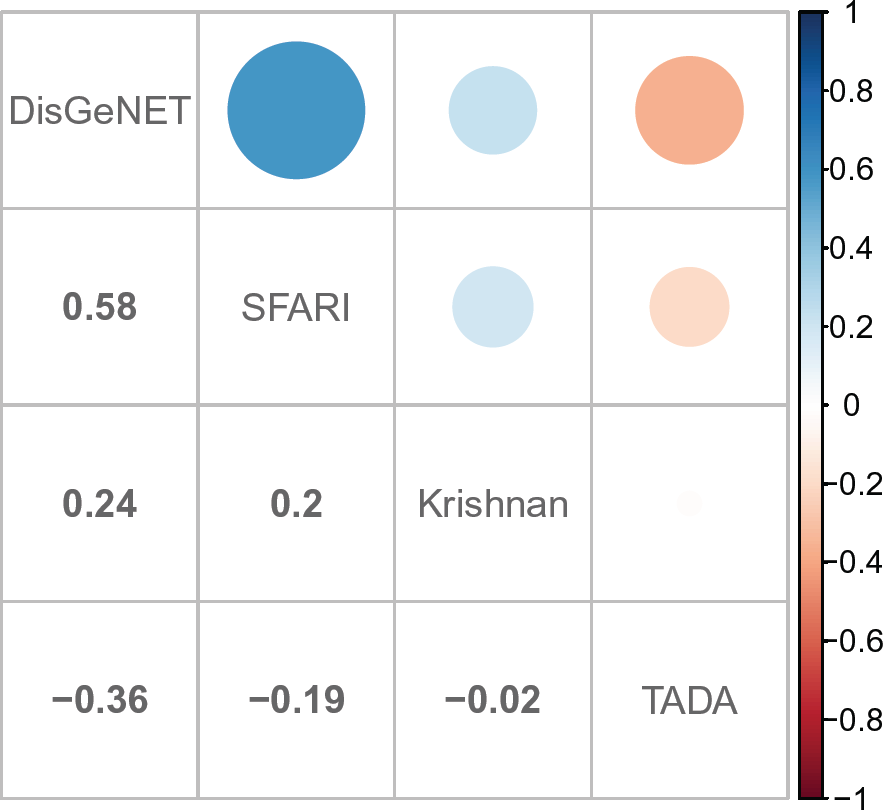
Pairwise correlation between the different ASD scoring systems studied. The size and colour of the circles correspond to the magnitude and sign of the correlation, respectively

Table 3 shows the correlation found between each of the scoring systems and the mean level of expression of the genes. Parallel to the results found above, The SFARI, Krishnan and DisGeNET scores have positive correlations, while Sanders TADA score appears to be independent.

**Table 3.** Correlation between different ASD scoring systems and the mean level of expression of the genes

|  | SFARI | Krishnan | DisGeNET | TADA |
| --- | --- | --- | --- | --- |
| Correlation | 0.35 | 0.35 | 0.19 | -0.01 |
| p-value | $4 \times 10^{-17}$ | 0 | 0.003 | 0.097 |

#### 3.5.2. Relation between mean expression and other neuronal disorders

The gene scores for other neuronal disorders were obtained from DisGeNET. The disorders selected were Schizophrenia (Scz), Bipolar Disorder (BD), Intellectual Disability (ID), Depressive Disorder (DD) and Chronic Alcohol Intoxication (CAI).

A big proportion of the genes associated to all of these disorders belong to the SFARI Genes, as Table 4 shows, the highest being Intellectual Disability with 24% and the lowest Schizophrenia, with 18%.

**Table 4.** Number of genes associated to different neuronal disorders according to DisGeNET and percentage of genes that belong to the SFARI Genes list

|  | ASD | Scz | BD | ID | DD | CAI |
| --- | --- | --- | --- | --- | --- | --- |
| Total | 231 | 765 | 415 | 425 | 254 | 228 |
| % of SFARI | 61.9% | 18.0% | 22.2% | 24.2% | 21.7% | 18.9% |

Studying the scores associated to each of the disorders, Figure 11 shows that SFARI genes are not only over-represented in all disorders, but they also have higher scores than the rest of the genes associated to each disorder. This difference is statistically significant for all disorders except for Chronic Alcohol Intoxication.

**Fig. 11.**
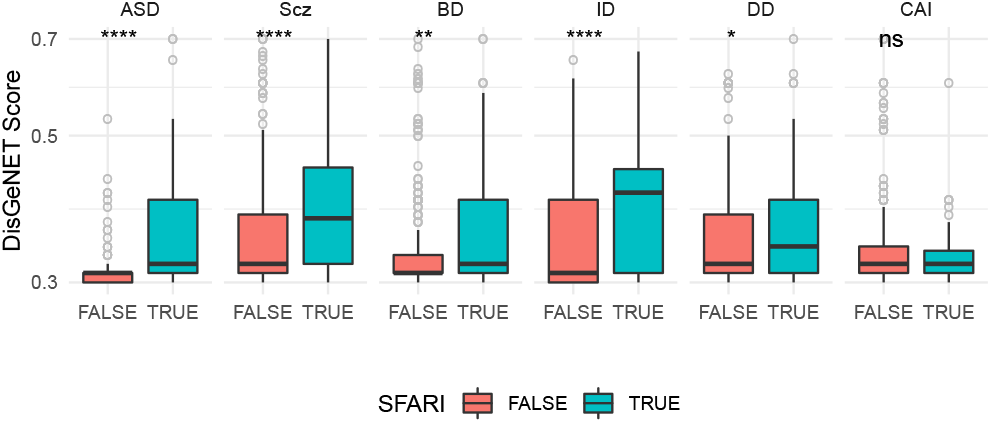
Box plot with the different genes associated to each disorder and their corresponding score, separating the SFARI genes from the rest. The asterisks at the top indicate how statistically significant is the difference between the two groups of genes.

Finally, calculating the correlation between the different scores and the mean expression of the genes, Table 5 shows ASD is the disorder with the highest correlation, followed by Schizophrenia and Bipolar Disorder, all three of them with p-values lower tan 0.05, but this relation weakens significantly when we remove the SFARI genes, as Table 6 shows, where the only disorder with a significant p-value is Schizophrenia, with a correlation of 0.09.

**Table 5.**
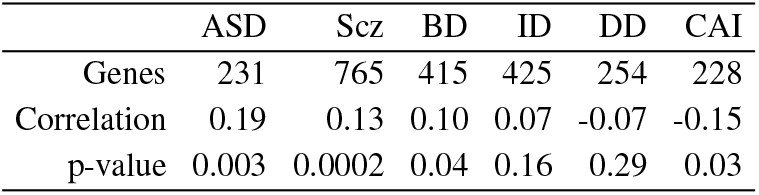
Correlation between the scores associated to different disorders by DisGeNET and the mean level of expression of the genes

|  | ASD | Scz | BD | ID | DD | CAI |
| --- | --- | --- | --- | --- | --- | --- |
| Genes | 231 | 765 | 415 | 425 | 254 | 228 |
| Correlation | 0.19 | 0.13 | 0.10 | 0.07 | -0.07 | -0.15 |
| p-value | 0.003 | 0.0002 | 0.04 | 0.16 | 0.29 | 0.03 |

**Table 6.** Correlation between the scores associated to different disorders by DisGeNET, removing the SFARI genes, and the mean level of expression of the genes

|  | ASD | Scz | BD | ID | DD | CAI |
| --- | --- | --- | --- | --- | --- | --- |
| Non-SFARI genes | 88 | 627 | 323 | 322 | 199 | 185 |
| Correlation | 0.02 | 0.09 | 0.07 | -0.06 | -0.08 | -0.09 |
| p.value | 0.86 | 0.02 | 0.24 | 0.31 | 0.24 | 0.20 |

Taken together these results demonstrate that the unexpected profile of the mean level of expression observed for genes in the SFARI-gene list is not related to the method of scoring used by SFARI as evidenced by it’s persistence in other independent scoring systems. Additionally, we show that this feature is not restricted to ASD, but found also in other neurological disorders including Schizophrenia, Bipolar Disorder, Intellectual Disability and Depressive Disorder despite them having far fewer SFARI genes present in their disease associated gene lists.

## 4 Discussion

SFARI genes have a lower percentage of differentially expressed genes than non-SFARI genes with neuronal function, as well as a lower log fold change magnitude than all non-SFARI genes, regardless of neuronal function. When separating the SFARI genes by score, we find that the higher the SFARI score, the lower the log fold change magnitude of the genes. This decrease within SFARI scores and between SFARI genes and other genes is partly explained by the high mean level of expression found for SFARI genes. However, even after correcting for this effect a small bias remains; we can only conclude this is likely due to other as yet unknown biological factors.

Modules derived from our gene co-expression network showed no significant correlation between the module diagnosis status and module enrichment for SFARI genes. This suggests that even though SFARI genes do cluster together within modules, these modules are not especially disrupted by ASD. The bias gene expression level in modules was unexpected, since the network was built using pairwise gene correlations, and the correlation metric is invariant to linear transformations. This suggests there may be more factors involved in this bias, and the level of expression may only be a confounding factor for another underlying trait.

Contrary to the results observed at *gene-level* and *module-level*, we demonstrate that SFARI-gene status can be successfully used in combination with differential gene expression data when considered at the *systems-level*. This suggests that local information is not sufficient to describe the complex role SFARI genes play in gene-expression profiles and their dysregulation in ASD, but instead requires the whole network to model this intricate system.

The classifier used here was chosen for its explicit interpretability rather than predictive power per se, so it would be interesting in the future to determine whether different classification approaches are able to further improve on classification performance and to what extent this approach can generalise to other biological settings. Models could further be developed to embrace a semi-supervised learning approach. SFARI genes are confirmed disease genes, so it is valid to label them as positive, but the opposite is not true for non-SFARI genes since we do not know whether they are associated with ASD or not. Instead of labelling them as strictly negative, a better approach might be to leave them unlabelled, as the PU Learning methodology proposes (Li and Liu, 2005), and which has already been used for disease gene identification in protein-protein interaction networks with reported good performance (Yang *et al*., 2012). We also consider that the selection of which features to extract and use from the co-expression network warrants further investigation since much of the information about the structure of the network is lost. Using a classifier directly on the network, as reported elsewhere (Di Nanni *et al*., 2019), could be productive in further optimising classification performance.

The relationship found between SFARI genes and the mean level of expression was significant and persisted throughout all of the levels of our analysis. Although we don’t know what could be causing this, a possible explanation for it, as well as for the bias within the SFARI scores, could be a bias in the selection of the participants for genetic experiments; focusing mostly on people with moderate to severe ASD and overlooking people with milder cases, since (Chang et al., 2014) found that the severity of ASD phenotype is directly related to expression level of the genes, but since no information about the severity of the ASD of the participants is in the Spark Gene List (Feliciano *et al*., 2019), on which the SFARI-gene selection and scoring criteria rely is available we cannot assess this possibility.

Importantly, the bias found in the mean expression of SFARI genes is also present in several other ASD gene scoring systems with the exception of the TADA-score. This observation could be an indirect effect of the incorporation of SFARI-gene related information into the generation of DisGeNET and Krishnan scoring systems, but it is not obvious based on how the very different methods by which the scores are calculated how this would result in such a strong effect. Similarly, when we look at the DisGeNet scores of SFARI genes for other neuronal disorders, they have statistically significantly higher scores than the rest of the genes associated with each disorder. This raises the intriguing possibility that there may be significant shared molecular aetiology between these neurological diseases.

## 5 Conclusion

The relationship between SFARI genes and gene expression data is subtle and complicated needing information derived from the whole gene coexpression network to be modelled accurately. The use of SFARI-gene lists to validate results from differential expression analysis or from modules derived from gene co-expression networks even when they have a high module correlation with diagnosis status is not appropriate, yet widely done. We have shown that these patterns are not significantly associated with SFARI genes. Rather, careful *systems-level* network analysis and the use of machine learning models to combine different sources of data in disease settings can prove to be highly effective at least for the novel candidate gene prediction approach addressed here. We also emphasise the importance of carefully studying the innate features of the gene expression data used in any given study as exemplified by the sizeable gene expression level feature found for SFARI genes which, to our knowledge, has been overlooked until now. Understanding the intricate behaviour of SFARI genes is crucial, as their influence permeates many other ASD scoring systems, and even impacts data from other neurodevelopmental disorders. Further studies into the origins of this observed gene expression level bias and its origins will undoubtedly help us better understand ASD in the future.

## References

Araujo, D., Toriumi, K., Ochoa, C., Kulkarni, A., Anderson, A., Harper, M., Usui, N., Ellegood, J., Lerch, J., Birnbaum, S., Tucker, H., Powell, C., and Konopka, G. (2017). Foxp1 in forebrain pyramidal neurons controls gene expression required for spatial learning and synaptic plasticity. The Journal of Neuroscience, 37, 1005–17.

Ashburner, M., Ball, C., Blake, J., Botstein, D., Butler, H., and Cherry, J. (2000). Gene ontology: Tool for the unification of biology. The Gene Ontology Consortium. Nat Genet, 25, 25–29.

Banerjee-Basu, S. and Packer, A. (2010). Sfari gene: an evolving database for the autism research community. Disease Models & Mechanisms, 3(3-4), 133–135.

Berto, S., Wang, G., Germi, J., Lega, B., and Konopka, G. (2018). Human genomic signatures of brain oscillations during memory encoding. Cerebral Cortex, 28(5), 1733–1748. Copyright: This record is sourced from MEDLINE/PubMed, a database of the U.S. National Library of Medicine.

Brueggeman, L., Koomar, T., and Michaelson, J. (2020). Forecasting risk gene discovery in autism with machine learning and genome-scale data. Scientific Reports, 10, 4569.

Chang, J., Gilman, S., Chiang, A., Sanders, S., and Vitkup, D. (2014). Genotype to phenotype relationships in autism spectrum disorders. Nature neuroscience, 18.

Chawla, N. V., Bowyer, K. W., Hall, L. O., and Kegelmeyer, W. P. (2002). Smote: Synthetic minority over-sampling technique. J. Artif. Int. Res., 16(1), 321–357.

Cogill, S. and Wang, L. (2016). Support vector machine model of developmental brain gene expression data for prioritization of Autism risk gene candidates. Bioinformatics, 32(23), 3611–3618.

Consortium, T. and Acencio, M. (2018). The gene ontology resource: 20 years and still going strong. Nucleic Acids Research, 49, gky1055.

Croft, D., Fabregat Mundo, A., Haw, R., Milacic, M., Weiser, J., Wu, G., Caudy, M., Garapati, P., Gillespie, M., Kamdar, M., Jassal, B., Jupe, S., Matthews, L., May, B., Palatnik, S., Rothfels, K., Shamovsky, V., Song, H., Williams, M., and D’Eustachio, P. (2013). The reactome pathway knowledgebase. Nucleic acids research, 42.

Di Nanni, N., Bersanelli, M., Cupaioli, F.A., Milanesi, L., Mezzelani, A., and Mosca, E. (2019). Network-based integrative analysis of genomics, epigenomics and transcriptomics in autism spectrum disorders. International Journal of Molecular Sciences, 20(13).

Durinck, S., Moreau, Y., Kasprzyk, A., Davis, S., De Moor, B., Brazma, A., and Huber, W. (2005). Biomart and bioconductor: a powerful link between biological databases and microarray data analysis. Bioinformatics, 21, 3439–3440.

Edgar, R. and Lash, A. (2002). 6. the gene expression omnibus (geo): A gene expression and hybridization repository. Nucleic Acids Res.

Feliciano, P., Zhou, X., Astrovskaya, I., Turner, T., Tianyun, W., Brueggeman, L., Barnard, R., Hsieh, A., Green Snyder, L., Muzny, D., Sabo, A., Gibbs, R., Eichler, E., O’Roak, B., Michaelson, J., Volfovsky, N., Shen, Y., and Chung, W. (2019). Exome sequencing of 457 autism families recruited online provides evidence for autism risk genes. npj Genomic Medicine, 4.

Gandal, M. J., Haney, J. R., Parikshak, N. N., Leppa, V., Ramaswami, G., Hartl, C., Schork, A. J., Appadurai, V., Buil, A., Werge, T. M., Liu, C., White, K. P., Horvath, S., and Geschwind, D. H. (2018). Shared molecular neuropathology across major psychiatric disorders parallels polygenic overlap. Science, 359(6376), 693–697.

Gaugler, T., Klei, L., Sanders, S., Bodea, C., Goldberg, A., Lee, A., Mahajan, M., Manaa, D., Pawitan, Y., Reichert, J., Ripke, S., Sandin, S., Sklar, P., Svantesson, O., Reichenberg, A., Hultman, C. M., Devlin, B., Roeder, K., and Buxbaum, J. (2014). Most genetic risk for autism resides with common variation. Nature genetics, 46.

Gokoolparsadh, A., Fang, Z., Braidy, N., and Voineagu, I. (2017). Topoisomerase i inhibition leads to length-dependent gene expression changes in human primary astrocytes. Genomics Data, 11, 113–115.

Gupta, S., Ellis, S., Ashar, F., Moes, A., Bader, J., Zhan, J., West, A., and Arking, D. (2014). Transcriptome analysis reveals dysregulation of innate immune response genes and neuronal activity-dependent genes in autism. Nature communications, 5, 5748.

Haslinger, D., Waltes, R., Yousaf, A., Lindlar, S., Schneider, I., Lim, C., Tsai, M.-M., Garvalov, B., Acker-Palmer, A., Krezdorn, N., Rotter, B., Acker, T., Guillemin, G., Fulda, S., Freitag, C., and Chiocchetti, A. (2018). Loss of the chr16p11.2 asd candidate gene qprt leads to aberrant neuronal differentiation in the sh-sy5y neuronal cell model. Molecular Autism, 9.

He, X., Sanders, S., Liu, l., De Rubeis, S., Lim, E., Sutcliffe, J., Schellenberg, G., Gibbs, R., Daly, M., Buxbaum, J., State, M., Devlin, B., and Roeder, K. (2013). Integrated model of de novo and inherited genetic variants yields greater power to identify risk genes. PLoS genetics, 9, e1003671.

Hoerl, A. E. and Kennard, R. W. (2000). Ridge regression: Biased estimation for nonorthogonal problems. Technometrics, 42(1), 80–86.

Hu, V. and Devlin, D. (2019). Asd phenotype—genotype associations in concordant and discordant monozygotic and dizygotic twins stratified by severity of autistic traits. International Journal of Molecular Sciences, 20, 3804.

Iossifov, I., O’Roak, B., Sanders, S., Ronemus, M., Krumm, N., Levy, D., Stessman, H., Witherspoon, K., Vives, L., Patterson, K., Smith, J., Paeper, B., Nickerson, D., Dea, J., Dong, S., Gonzalez, L., Mandell, J., Mane, S., Murtha, M., and Wigler, M. (2014). The contribution of de novo coding mutations to autism spectrum disorder. Nature, 515.

Jiang, H. and Nachum, O. (2019). Identifying and correcting label bias in machine learning.

Kanehisa, M. and Goto, S. (2000). Kegg: kyoto encyclopedia of genes and genomes. Nucleic acids research, 28, 27–30.

Khatri, N. and Man, H.-Y. (2019). The autism and angelman syndrome protein ube3a/e6ap: The gene, e3 ligase ubiquitination targets and neurobiological functions. Frontiers in Molecular Neuroscience, 12.

Krishnan, A., Zhang, R., Yao, V., Theesfeld, C., Wong, A., Tadych, A., Volfovsky, N., Packer, A., Lash, A., and Troyanskaya, O. (2016). Genome-wide prediction and functional characterization of the genetic basis of autism spectrum disorder. Nature neuroscience, 19.

Langfelder, P. and Horvath, S. (2012). Fast R functions for robust correlations and hierarchical clustering. Journal of Statistical Software, 46(11), 1–17.

Leek, J. T., Johnson, W. E., Parker, H. S., Fertig, E. J., Jaffe, A. E., Storey, J. D., Zhang, Y., and Torres, L. C. (2019). sva: Surrogate Variable Analysis. R package version 3.32.1.

Li, X.-L. and Liu, B. (2005). Learning from positive and unlabeled examples with different data distributions. In Proceedings of the 16th European Conference on Machine Learning, ECML’05, page 218–229, Berlin, Heidelberg. Springer-Verlag.

Lin, Y., Afshar, S., Rajadhyaksha, A. M., Potash, J. B., and Han, S. (2020). A machine learning approach to predicting autism risk genes: Validation of known genes and discovery of new candidates. Frontiers in Genetics, 11, 1051.

Liu, X., Han, D., Somel, M., Jiang, X., Hu, H., Guijarro, P., Zhang, N., Mitchell, A., Halene, T., Ely, J., Sherwood, C., Hof, P., Qiu, Z., Pääbo, S., Akbarian, S., and Khaitovich, P. (2016). Disruption of an evolutionarily novel synaptic expression pattern in autism. PLOS Biol, 14.

Lombardo, M., Moon, H., Su, J., Palmer, T., Courchesne, E., and Pramparo, T. (2017). Maternal immune activation dysregulation of the fetal brain transcriptome and relevance to the pathophysiology of autism spectrum disorder. Molecular Psychiatry, 23.

Lord, C., Rutter, M., and Le Couteur, A. (1994). Autism Diagnostic Interview-Revised: A revised version of a diagnostic interview for caregivers of individuals with possible pervasive developmental disorders. Journal of Autism and Developmental Disorders, 24(5), 659–685.

Love, M. I., Huber, W., and Anders, S. (2014). Moderated estimation of fold change and dispersion for rna-seq data with deseq2. Genome Biology, 15, 550.

Marin, I. (2010). Animal hect ubiquitin ligases: Evolution and functional implications. BMC evolutionary biology, 10, 56.

Martin, J., Cooper, M., Hamshere, M., Pocklington, A., Scherer, S., Kent, L., Gill, M., Owen, M., Williams, N., O’Donovan, M., Thapar, A., and Holmans, P. (2014). Biological overlap of attention-deficit/hyperactivity disorder and autism spectrum disorder: Evidence from copy number variants. Journal of the American Academy of Child & Adolescent Psychiatry, 53.

Nowakowski, T., Bhaduri, A., Pollen, A., Alvarado, B., Mostajo Radji, M., Di Lullo, E., Haeussler, M., Sandoval-Espinosa, C., Liu, J., Velmeshev, D., Ounadjela, J., Shuga, J., Wang, X., Lim, D., West, J., Leyrat, A., Kent, W., and Kriegstein, A. (2017). Spatiotemporal gene expression trajectories reveal developmental hierarchies of the human cortex. Science, 358, 1318–1323.

Pinto, D., Delaby, E., Merico, D., Barbosa, M., Merikangas, A., Klei, L., Thiruvahindrapuram, B., Xu, X., Ziman, R., Wang, Z., Vorstman, J. A. S., Thompson, A., Regan, R., Pilorge, M., Pellecchia, G., Pagnamenta, A. T., Oliveira, B., Marshall, C. R., Magalhaes, T. R., Lowe, J. K., Howe, J. L., Griswold, A. J., Gilbert, J., Duketis, E., Dombroski, B.A., De Jonge, M.V., Cuccaro, M., Crawford, E. L., Correia, C. T., Conroy, J., Conceição, I. C., Chiocchetti, A. G., Casey, J. P., Cai, G., Cabrol, C., Bolshakova, N., Bacchelli, E., Anney, R., Gallinger, S., Cotterchio, M., Casey, G., Zwaigenbaum, L., Wittemeyer, K., Wing, K., Wallace, S., van Engeland, H., Tryfon, A., Thomson, S., Soorya, L., Rogé, B., Roberts, W., Poustka, F., Mouga, S., Minshew, N., McInnes, L. A., McGrew, S. G., Lord, C., Leboyer, M., Le Couteur, A. S., Kolevzon, A., Jiménez González, P., Jacob, S., Holt, R., Guter, S., Green, J., Green, A., Gillberg, C., Fernandez, B. A., Duque, F., Delorme, R., Dawson, G., Chaste, P., Café, C., Brennan, S., Bourgeron, T., Bolton, P. F., Bölte, S., Bernier, R., Baird, G., Bailey, A. J., Anagnostou, E., Almeida, J., Wijsman, E. M., Vieland, V. J., Vicente, A. M., Schellenberg, G. D., Pericak-Vance, M., Paterson, A. D., Parr, J. R., Oliveira, G., Nurnberger, J. I., Monaco, A. P., Maestrini, E., Klauck, S. M., Hakonarson, H., Haines, J. L., Geschwind, D. H., Freitag, C. M., Folstein, S. E., Ennis, S., Coon, H., Battaglia, A., Szatmari, P., Sutcliffe, J. S., Hallmayer, J., Gill, M., Cook, E. H., Buxbaum, J. D., Devlin, B., Gallagher, L., Betancur, C., and Scherer, S. W. (2014). Convergence of genes and cellular pathways dysregulated in autism spectrum disorders. American Journal of Human Genetics, 94(5), 677–694.

Piñero, J., Ramírez-Anguita, J., Saüch-Pitarch, J., Ronzano, F., Centeno, E., Sanz, F., and Furlong, L. I. (2019). The disgenet knowledge platform for disease genomics: 2019 update. Nucleic acids research, 48.

Quesnel-Vallières, M., Weatheritt, R., Cordes, S., and Blencowe, B. (2018). Autism spectrum disorder: insights into convergent mechanisms from transcriptomics. Nature Reviews Genetics, 20.

Retuerto, A., Cantor, R., Gleeson, J., Ustaszewska, A., Schackwitz, W., Pennacchio, L., and Geschwind, D. (2008). Association of common variants in the joubert syndrome gene (ahi1) with autism. Human molecular genetics, 17, 3887–96.

Sayers, E., Barrett, T., Benson, D., Bolton, E., Bryant, S., Canese, K., Chetvernin, V., Church, D., Dicuccio, M., Federhen, S., Feolo, M., Fingerman, I., Geer, L., Helmberg, W., Kapustin, Y., Krasnov, S., Landsman, D., Lipman, D., lu, Z., and Ye, J. (2011). Database resources of the national center for biotechnology information. Nucleic acids research, 40, D13–25.

Schriml, L., Mitraka, E., Munro, J., Tauber, B., Schor, M., Nickle, L., Felix, V., Jeng, L., Bearer, C., Lichenstein, R., Bisordi, K., Campion, N., Hyman, B., Kurland, D., Oates, C., Kibbey, S., Sreekumar, P., Le, C., Giglio, M., and Greene, C. (2018). Human disease ontology 2018 update: classification, content and workflow expansion. Nucleic acids research, 47.

Suetterlin, P., Hurley, S., Mohan, C., Riegman, K., Pagani, M., Caruso, A., Ellegood, J., Galbusera, A., Crespo Enriquez, I., Michetti, C., Yee, Y., Ellingford, R., Brock, O., Delogu, A., Francis-West, P., Lerch, J., Scattoni, M. L., Gozzi, A., Fernandes, C., and Basson, M. (2018). Altered neocortical gene expression, brain overgrowth and functional over-connectivity in chd8 haploinsufficient mice. Cerebral Cortex, 28.

Tangsuwansri, C., Saeliw, T., Thongkorn, S., Chonchaiya, W., Suphapeetiporn, K., Mutirangura, A., Tencomnao, T., Hu, V. W., and Sarachana, T. (2018). Investigation of epigenetic regulatory networks associated with autism spectrum disorder (asd) by integrated global line-1 methylation and gene expression profiling analyses. PLOS ONE, 13(7), 1–27.

Walker, S. and Scherer, S. (2013). Identification of candidate intergenic risk loci in autism spectrum disorder. BMC genomics, 14, 499.

Wang, P., Zhao, D., Lachman, H., and Zheng, D. (2018). Enriched expression of genes associated with autism spectrum disorders in human inhibitory neurons. Translational Psychiatry, 8.

Wright, C., Shin, J., Rajpurohit, A., Deep-Soboslay, A., Collado-Torres, L., Brandon, N., Hyde, T., Kleinman, J., Jaffe, A., Cross, A., andWeinberger, D. (2017). Altered expression of histamine signaling genes in autism spectrum disorder. Translational Psychiatry, 7, e1126.

Yang, P., Li, X.-L., Mei, J.-P., Kwoh, C.-K., andNg, S.-K. (2012). Positive-unlabeled learning for disease gene identification. Bioinformatics, 28(20), 2640–2647.

Yu, G., Wang, L.-G., Han, Y., and He, Q.-Y. (2012). clusterprofiler: an r package for comparing biological themes among gene clusters. OMICS: A Journal of Integrative Biology, 16(5), 284–287.

Yu, Q. and He, Z. (2017). Comprehensive investigation of temporal and autism-associated cell type composition-dependent and independent gene expression changes in human brains open. Scientific Reports, 7.

Zoubarev, A., Hamer, K. M., Keshav, K. D., McCarthy, E. L., Santos, J. R. C., Van Rossum, T., McDonald, C., Hall, A., Wan, X., Lim, R., Gillis, J., and Pavlidis, P. (2012). Gemma: are source for the reuse, sharing andmeta-analysis of expression profiling data. Bioinformatics, 28(17), 2272–2273.

